# Dynamical reorganization of transcriptional events governs robust Nanog heterogeneity

**DOI:** 10.1101/527747

**Authors:** Tagari Samanta, Sandip Kar

**Affiliations:** Department of Chemistry, IIT Bombay, Powai, Mumbai - 400076, India

## Abstract

Nanog maintains pluripotency of embryonic stem cells (ESC’s), while demonstrating high expression heterogeneity within an ESC population. Intriguingly, in ESC’s, the overall heterogeneity at the Nanog mRNA level under various culture conditions gets precisely partitioned into intrinsic (~45%) and extrinsic (~55%) fluctuations. However, the dynamical origin of such a robust transcriptional noise regulation, still remains illusive. Herein, we conceived a new stochastic simulation strategy centered around Gillespie’s stochastic simulation algorithm to efficiently capture fluctuations of different origins that are operative within a simple Nanog transcriptional regulatory network. Our model simulations reconcile the strict apportioning of Nanog transcriptional fluctuation, while predicting possible experimental scenarios to avoid such an exact noise segregation. Importantly, model analyses reveal that different culture conditions essentially preserve the Nanog expression heterogeneity by altering the dynamics of transcriptional events. These insights will be essential to systematically maneuver cell-fate decision making events of ESC’s for therapeutic applications.

## Introduction

Embryonic stem cells (ESCs) have a unique ability to either maintain their stemness via self-renewal or differentiate into a particular cell-type under specific growth conditions (Hyslop et al., 2005; Lie et al., 2012; Young, 2011). Thus, ESC’s will be highly relevant for developing stem cell based treatment strategies, provided, we figure out how to tune the specific cell fate determining molecular regulators of ESC’s by altering the culture conditions precisely. Among these molecular regulators, Nanog (a transcription factor) plays a decisive role (Ambrosetti et al., 2000; Boyer et al., 2006; Loh et al., 2006; Macarthur et al., 2013; Mitsui et al., 2003; Niwa, 2007). A higher level of Nanog expression maintains ESC’s in a pluripotent like state, whereas ESC’s expressing low levels of Nanog are more prone to differentiate (Chambers et al., 2007; Fischer et al., 2010; Glauche et al., 2010; Kalmar et al., 2009; Singh et al., 2007). Intriguingly, Nanog displays a highly heterogeneous expression pattern within a population of ESC’s (Eldar and Elowitz, 2010; Navarro et al., 2012). It was argued qualitatively that fluctuations in different form present in and around the ESC’s produce such heterogeneity in Nanog protein expression (Faddah et al., 2013; Filipczyk et al., 2013; Kalmar et al., 2009), which in turn affects the stemness/differentiation balance of ESC’s. Unfortunately, how exactly fluctuations maintain Nanog expression heterogeneity in ESC’s, still remains unclear. It is thus imperative to investigate how fluctuations, at the first place, influence the Nanog transcriptional events to orchestrate the Nanog expression heterogeneity.

Deciphering the impact of fluctuations on Nanog transcription process definitely requires a precise quantification of the effect of extrinsic (due to cell to cell variability within a cellular population) and intrinsic (molecular fluctuation inherent to the gene expression process) fluctuations on Nanog mRNA expression. Recently, Ochiai et al. (Ochiai et al., 2014) achieved this feat experimentally by performing allele-specific single molecule RNA fluorescent in situ hybridization (smFISH) experiments in a reporter cell line (NMP cell line), and quantified the contributions of different noise sources in Nanog mRNA regulation by following the classical work of Elowitz et al. (Elowitz et al., 2002). Interestingly, they demonstrated that the overall fluctuation at the Nanog transcription level is robustly partitioned between intrinsic and extrinsic fluctuations by conserving a ratio of ~ 45:55 under a variety of culture conditions consisting of either simple serum, or only PD0325901 (MEK inhibitor) inhibitor, and a range of 2i (PD0325901 and CHIR99021 (GSK3 inhibitor)) conditions. How and why this stringent noise partitioning ratio is achieved every time under diverse culture conditions? Moreover, how to bypass such robust splitting of intrinsic and extrinsic fluctuations in Nanog transcription? Answering these questions are not straightforward, and need improved quantitative modeling framework to precisely disentangle the effect of fluctuations in the Nanog transcription.

In literature, mathematical and computational modeling studies had been employed quite frequently to obtain crucial insight related to transcriptional noise regulation in different biological contexts (McAdams and Arkin, 1997; Spudich and Koshland, 1976). Unfortunately, in many of these studies, it is hard to accurately segregate the contributions of the intrinsic and extrinsic fluctuations quantitatively. Often fluctuations due to extrinsic factors are underrepresented due to unavailability of appropriate modeling technique to incorporate them into the standard stochastic simulation frameworks such as Gillespie’s stochastic simulation algorithm (SSA) (Gillespie, 1976). In this article, we put forward a new generalized numerical recipe capable of accounting for different kinds of extrinsic fluctuations within a stochastic mathematical modeling framework. We employed this new technique to simulate a minimal gene regulatory network that governs the Nanog dynamics in ECS’s, and showed how and why a ~ 45:55 ratio of intrinsic and extrinsic fluctuations is maintained at the Nanog mRNA level. Not only that, our stochastic simulations predict how to avoid such rigid partitioning of noise regulation by altering culture conditions. Finally, we provided the rationale behind 45:55 ratio maintenance in Nanog transcription, which will be important in future to alter the balance between stemness and differentiation in ESC’s.

## The Model

We have considered a simple gene regulatory network of Nanog regulation (Fig. 1A) in ESC’s, where Nanog transcriptionally activates its positive regulators Oct4 and Sox2, and these two proteins forms a heterodimer that triggers Nanog transcription in a positive feedback manner (Ambrosetti et al., 2000; Boyer et al., 2006; Chickarmane et al., 2006; Festuccia et al., 2012; Mitsui et al., 2003; Niwa, 2007). The transcriptional network of Nanog has been modeled by keeping in the constructs introduced by Ochiai et al. (Ochiai et al., 2014) in the reporter NMP cell lines. We assumed that the promoter for the *Nanog* gene can either be in ON (Ga_N_) or OFF (Gi_N_) state (Fig. 1A). Ga_N_ will produce both the Nanog mRNA’s (M_MS2_ and M_PP7_) that are detectable through smFISH experiment (Chao et al., 2008; Dictenberg, 2012; Guet et al., 2008). Both of these transcripts of Nanog will yield the same Nanog protein (P_N_) that will eventually form a dimer (P2_N_) (Wang et al., 2008), and cause the transcriptional activation of *Oct4* and *Sox2* genes by bringing the corresponding promoters from OFF (Gi_O_ and Gi_S_) to ON (Ga_O_ and Ga_S_) states respectively. These activated promoters form the corresponding mRNA’s (M_O_ and M_S_) and proteins (P_O_ and P_S_) of Oct4 and Sox2. The heterodimer (OS) of P_O_ and P_S_ positively regulates the Nanog transcription by activating the Nanog promoter (Festuccia et al., 2012). The effect of serum induction has been included in all the promoter activation events as shown in Fig. 1A. The ordinary differential equations (Supplementary file 1A) depicting the overall network model, the corresponding reaction events for stochastic simulations (Supplementary file 1B), and the values for kinetic parameters (Supplementary file 1C) involved are given in the supplementary file 1. Most of our kinetic parameters are obtained from experimental (Ochiai et al., 2014; Sharova et al., 2009; Wei et al., 2007) literature (highlighted in Supplementary file 1C).

**Fig. 1.**
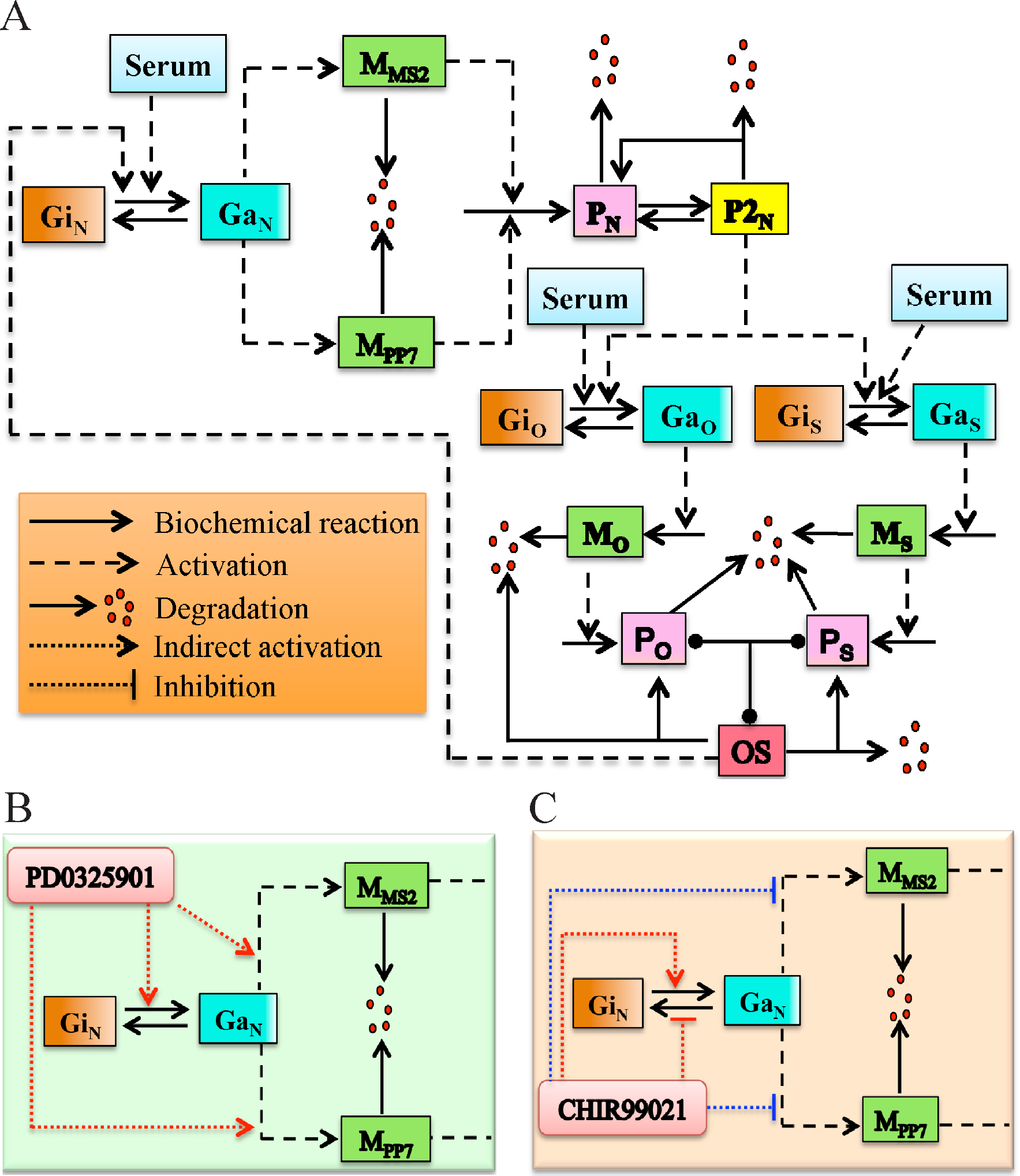
The proposed regulatory network of Nanog regulation in ESC’s under normal and inhibitory conditions. **(A)** The detailed network of the Nanog transcriptional regulation maintained by a positive feedback regulation from Oct4 and Sox2. The overall influence of the two inhibitors **(B)** PD0325901 and **(C)** CHIR99021 on Nanog transcriptional process are qualitatively shown. The detailed regulatory interactions are described in the text.

To simulate different culture conditions (serum and inhibitor conditions), we have systematically incorporated the effect of PD0325901 and CHIR99021 inhibitors in our model equations (Supplementary file 1A) in a phenomenological manner. PD0325901 inhibits the phosphorylation and activation of ERK signaling through the inhibition of MEK (Bain et al., 2007; Sebolt-Leopold and Herrera, 2004). Thus, PD0325901 increases Nanog expression by inhibiting ERK signaling (Fig. 1B), as activated form of ERK represses Nanog promoter activation (Figure supplement 1A) (Chen et al., 2015; Miyanari and Torres-Padilla, 2012). In our model, we have included this effect of PD0325901 inhibitor by considering a direct activation of Nanog promoter (Eq. 1, Supplementary file 1A) by PD0325901. At the same time, activated MEK indirectly represses Nanog transcription in mouse ES cells (Figure supplement 1A) (Hamazaki et al., 2006), which implies PD0325901 indirectly activates Nanog transcription (Fig. 1B). In Eq.2 and in Eq.3 (Supplementary file 1A), we have assumed a positive effect of PD0325901 on Nanog transcription phenomenologically, which is again happening due to two negative regulations (PD0325901 to MEK and MEK to Nanog transcription via some indirect path). We have chosen these phenomenological terms in Eq.1-3 (Supplementary file 1A) to obtain similar numbers of total Nanog mRNA molecule (Table 1), as reported by Ochiai. et. al. (Ochiai et al., 2014) under various doses of PD0325901 inhibitor (*I*_1_) concentrations.

In ESC’s, GSK-3 represses the Nanog promoter activation indirectly by activating Tcf3 (a transcriptional repressor for Nanog) via repressing β-catenin (Figure supplement 1B) (Godwin et al., 2017). Interestingly, it was shown recently that GSK3 also promotes Nanog promoter deactivation (Ochiai et al., 2014). This means that introducing CHIR99021 will activate the activation of Nanog promoter, and inhibit the deactivation process of the same via two different paths (Fig. 1C) (MURRAY et al., 2004; Storm et al., 2007; Ying et al., 2008). In our model (Eq.1), we have incorporated both these effects of CHIR99021 in Nanog promoter regulation phenomenologically. Additionally, we have introduced a negative influence of the CHIR99021 inhibitor in Nanog mRNA transcription (Eq.2 and Eq.3, Supplementary file 1A). This was required to reproduce the same number of Nanog mRNA molecules under different 2i conditions as reported in literature (Ochiai et al., 2014).

## Method

The deterministic model (Supplementary file 1A) proposed for the network in Fig. 1A is in terms of mass action kinetics, and it can readily be used for stochastic simulation (Supplementary file 1B) by employing SSA (Gillespie, 1976). The influence of intrinsic and extrinsic fluctuations present in the network can be calculated by using theory of covariance after obtaining the number of two different Nanog transcripts for about ~ 400 individual ESC’s (following experimental protocol) (Elowitz et al., 2002; Fu and Pachter, 2016). It turned out that our initial stochastic simulation (Figure supplement 2) using a traditional SSA predicts much higher influence of intrinsic fluctuations, and is far off from experimentally observed 45:55 ratio of intrinsic and extrinsic noise scenario. This result clearly reveals that simple SSA simulation of Nanog dynamics can not capture the extent of extrinsic variability’s that are present within an ESC’s population. This is due to the fact that in a simple SSA simulation framework, various extrinsic noise sources are not considered. For example, it is known experimentally that depending on the cell cycle phase of a particular cell, the promoter activation rate and the transcription rate varies quite a bit (Skinner et al., 2016; Zopf et al., 2013).

**Fig. 2.**
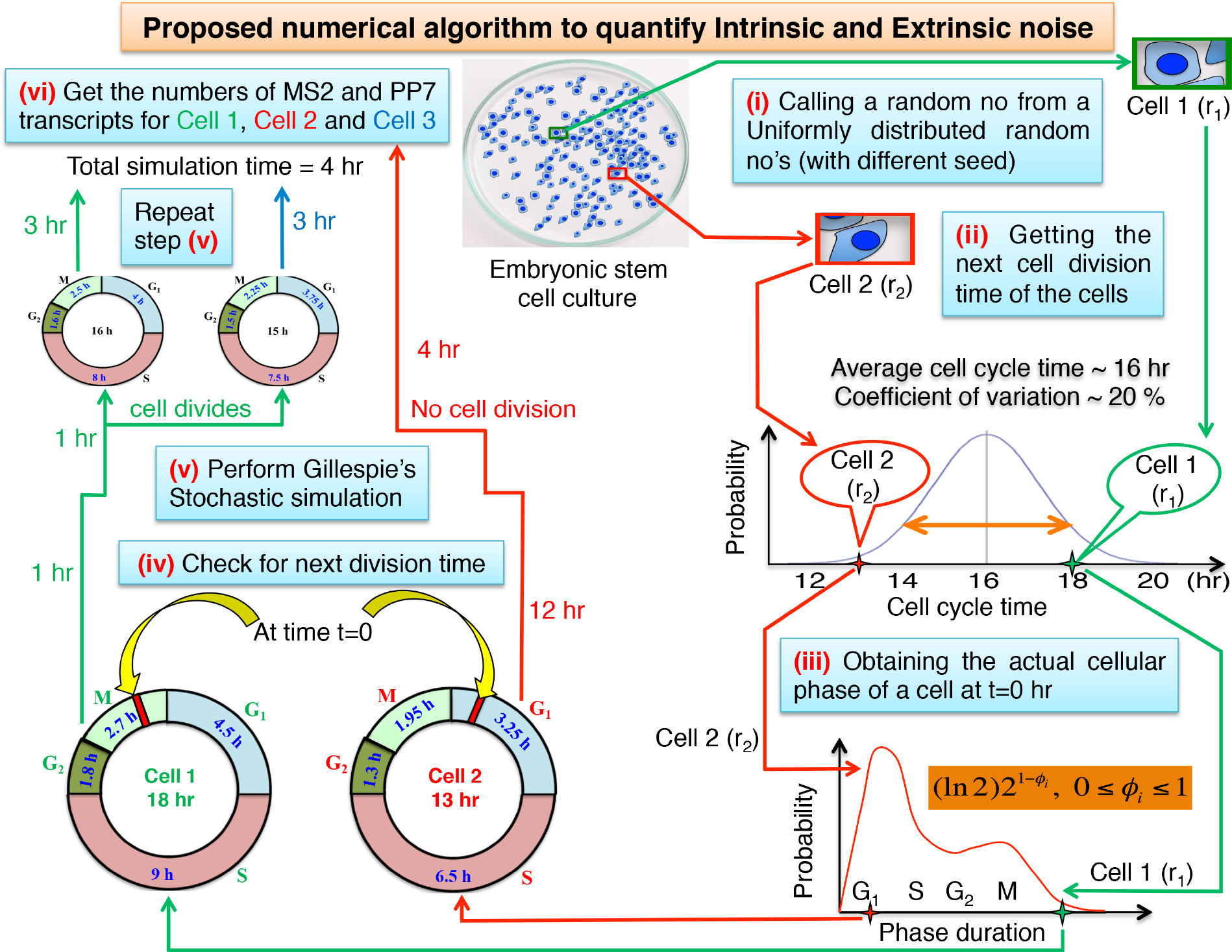
The proposed numerical recipe to capture extrinsic and intrinsic fluctuations accurately in Nanog transcriptional regulation. Two different scenarios’ i.e., stochastic simulation with (Cell 1) and without (Cell 2) cell division. The detailed steps of the numerical simulations are depicted in Figure supplement 3 schematically.

Since the experiments are performed with an asynchronous population of ESC’s, not all the cells are at the same phase of the cell cycle. Importantly, within the 4 hr experimental time frame (i.e., before performing the smFISH experiment at t = 4 hr after initiating experiment at t = 0 hr), some of the cells will definitely undergo cell division. This will be a big external perturbation, as all the network components will be divided among the two newborn cells, (Huh and Paulsson, 2011) and depending on the time of division and the natural time scale of the Nanog network, the system might or might not return back to its original dynamical state, when the smFISH experiment is performed. This may play a critical role in increasing the extrinsic fluctuations. How to incorporate all these extrinsic factors, especially the cell division phenomena, while performing our stochastic simulations?

To address this important question, we proposed a new numerical recipe to introduce the essential information we require from the cell division event without really considering a cell cycle model for ESC’s. It works in the following manner. For the time being, if we assume that the cell cycle dynamics of the ESC’s is happening independently of the Nanog network proposed in Fig. 1, then we require two crucial information’s while simulating the Nanog dynamics of a particular cell within an ESC’s population. At t = 0 hr, we need to know the next cell cycle time (like Cell 1, Fig. 2) of a particular ESC along with the current phase of cell cycle the cell is in at the same starting point of the stochastic simulation. This information’s will allow us to decide whether that particular ESC will divide or not within the 4 hr of time frame (before smFISH experiment), and also to set the transcription rates appropriately by knowing the actual cell cycle phase the cell is in (Figure supplement 3). Herein, to obtain the individual cell cycle period and the corresponding cell cycle phase that the ESC is in, we used two respective distributions as shown in Fig. 2 (for details see Figure supplement 3) in accordance with experimental findings (Hindley et al., 2013; Kar et al., 2009; Niepel et al., 2009; Skinner et al., 2016; White et al., 2005; Zopf et al., 2013). This provided us the crucial knowledge about which cell is going to divide and which one will not divide during the simulation, and we can further set the transcription and promoter activation rates accordingly before implementing SSA. We have described two representative scenarios’ that can appear during the stochastic simulations in Fig. 2.

## Results and discussion

### Contribution of different noise sources in Nanog mRNA expression variability

We implemented the algorithm (Fig. 2 and Figure supplement 3) to stochastically simulate the Nanog network (Fig. 1) to determine the noise contributions in the expression variability of Nanog mRNA. At first, we optimized the network in such a way that we could reproduce total number of Nanog mRNA under different culture conditions (serum and 2i condition, Table 1) as measured by Ochiai et al. (Ochiai et al., 2014). We calculated the noise contributions following the same procedure as implemented by Ochiai et al. (Ochiai et al., 2014)., while analyzing the smFISH data for 400 cells after 4 hrs of initial stimulation.

**Table 1.** Contribution of different noise sources calculated from stochastic simulation under different culture conditions (400 cells)

| Culture Conditions | PD0325901 ($\mu\text{M}$) | CHIR99021 ($\mu\text{M}$) | Mean mRNA | $\eta_{int}^2$ | $\eta_{ext}^2$ | $\eta_{tot}^2$ | % of intrinsic noise | % of extrinsic noise |
| --- | --- | --- | --- | --- | --- | --- | --- | --- |
| **Serum** | 0 | 0 | 42 | 0.522 | 0.667 | 1.19 | 44 | 56 |
| **PD0325901** | 0.2 | 0 | 103 | 0.294 | 0.341 | 0.635 | 46 | 54 |
|  | 0.5 | 0 | 217 | 0.229 | 0.272 | 0.508 | 46 | 54 |
|  | 0.75 | 0 | 293 | 0.21 | 0.199 | 0.408 | 52 | 48 |
|  | 1 | 0 | 323 | 0.166 | 0.202 | 0.369 | 45 | 55 |
| **CHIR99021** | 0 | 0.6 | 55 | 0.4 | 0.459 | 0.848 | 47 | 53 |
|  | 0 | 1.5 | 53 | 0.55 | 0.467 | 1.02 | 54 | 46 |
|  | 0 | 2.25 | 30 | 0.382 | 0.408 | 0.79 | 48 | 52 |
|  | 0 | 3 | 45 | 0.457 | 0.393 | 0.85 | 54 | 46 |
|  | 0 | 5 | 28 | 0.513 | 0.297 | 0.8 | 63 | 37 |
|  | 0 | 10 | 16 | 0.568 | 0.223 | 0.79 | 72 | 28 |
|  | 0 | 15 | 13 | 0.533 | 0.226 | 0.76 | 70 | 30 |
| **2i Condition** | 0.2 | 0.6 | 133 | 0.28 | 0.322 | 0.603 | 46 | 54 |
|  | 0.5 | 1.5 | 206 | 0.224 | 0.243 | 0.467 | 48 | 52 |
|  | 0.75 | 2.25 | 189 | 0.147 | 0.202 | 0.349 | 42 | 58 |
|  | 1 | 3 | 221 | 0.146 | 0.191 | 0.337 | 43 | 57 |
|  | 0.2 | 3 | 94 | 0.296 | 0.237 | 0.533 | 56 | 44 |
|  | 0.5 | 3 | 168 | 0.216 | 0.225 | 0.442 | 49 | 51 |
|  | 0.75 | 3 | 205 | 0.177 | 0.168 | 0.345 | 51 | 49 |
|  | 1 | 2 | 194 | 0.152 | 0.202 | 0.354 | 43 | 57 |
|  | 1 | 5 | 201 | 0.153 | 0.21 | 0.363 | 42 | 58 |
|  | 1 | 10 | 159 | 0.155 | 0.226 | 0.38 | 41 | 59 |

Our stochastic simulations showed a near 45:55 ratio of intrinsic and extrinsic fluctuation ratio under both serum and 2i culture conditions (Fig. 3A and Table 1). It is important to note that in our simulations, we have not been able to reproduce the absolute values of different noise levels, as observed experimentally, but our stochastic simulations capture the essential features and trends of the Nanog transcriptional noise regulation. We have performed noise calculation even for 2000 cells as well (Supplementary file 2), and the results remained consistent. Further, our simulations showed that intrinsic noise decreases with increasing the mean number of Nanog mRNA transcript (i.e., sum of MS2 and PP7 transcript) (Fig. 3B, Supplementary file 3), which is line with the experimental findings (Nicolas et al., 2017; Ochiai et al., 2014).

**Fig. 3.**
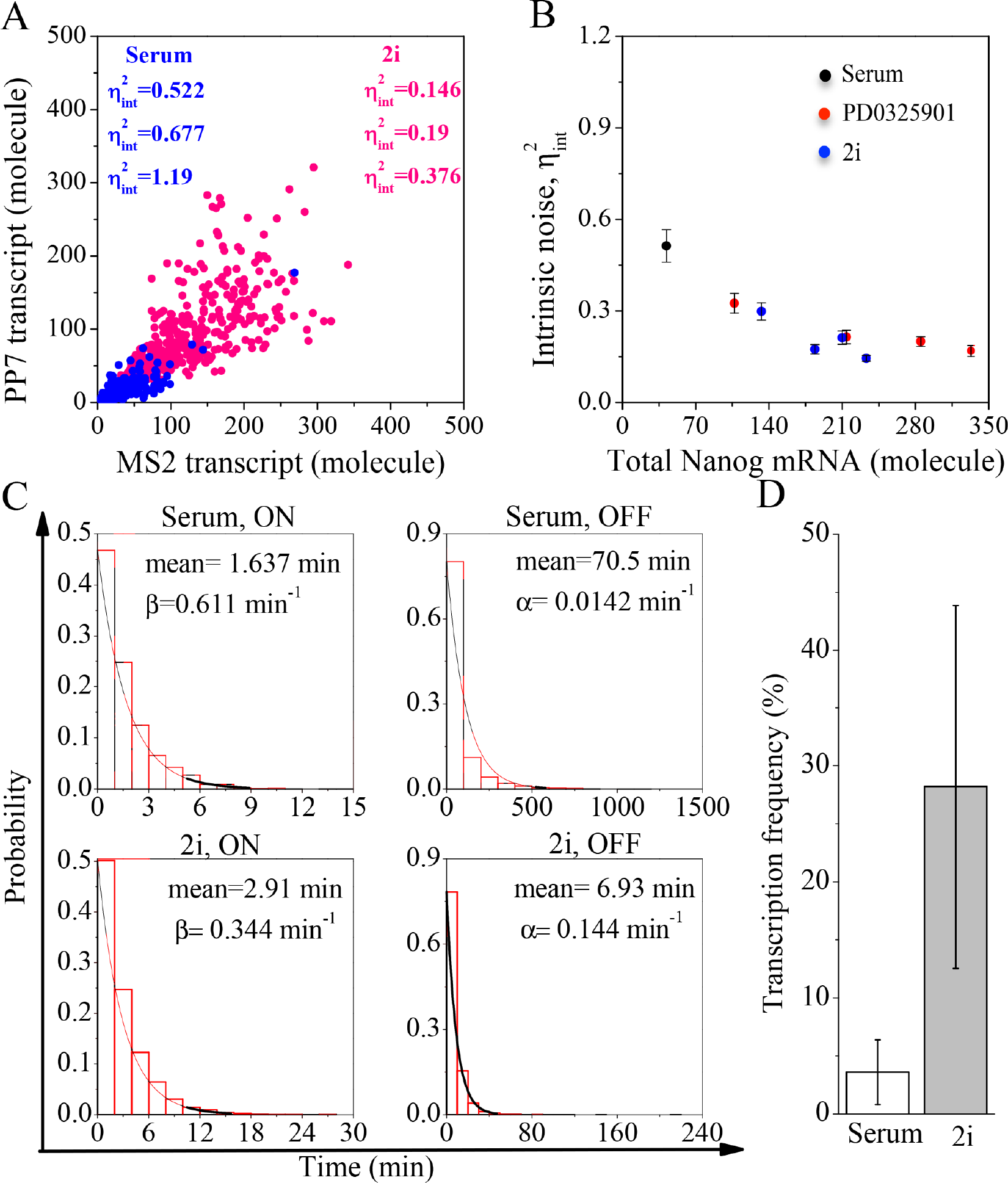
Model predicted Nanog transcriptional fluctuation pattern under different culture conditions. (**A)** Simulated scatter plot of Nanog mRNA transcripts (MS2 and PP7 tagged) under only serum and 2i culture conditions (for 400 cells). Each dot represents the number of Nanog-MS2 and Nanog-PP7 transcript in a single ESC. 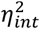, 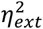 and 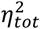 represent intrinsic, extrinsic and total noise, respectively. **(B)** Simulated intrinsic noise variation as a function of mean Nanog mRNA (for 400 cells) under different culture conditions (only serum, 2i and PD0325901 inhibitor condition). **(C)** ON and OFF time distributions of Nanog promoter under serum and 2i condition. Each probability distribution was fitted exponentially. **(D)** Distribution of transcription frequencies of Nanog cultured under serum and 2i conditions. We have observed 62 and 56 cells in serum and 2i culture condition respective over 4 hr to match experimental condition (Ochiai et al., 2014).

In literature, it has been demonstrated that promoter activation pattern significantly affects the transcriptional noise (Ochiai et al., 2014). Thus, we explored how the fluctuations at the Nanog transcriptional regulation is controlled by the promoter activity? Experimentally, it has been demonstrated that both the ON and OFF state durations of the promoter are quite different in the serum and 2i condition. Consistent with the experimental report, through our simulations, we got a longer mean duration of promoter being in the ON state under 2i condition (Fig. 3C, left lower panel) than in the serum condition (Fig. 3C, left upper panel). On the contrary, the mean duration of OFF time in serum condition (Fig. 3C, right upper panel) is much longer than that in 2i condition (Fig. 3C, right lower panel). Additionally, our stochastic simulations demonstrate that the mean transcriptional frequency under 2i condition (Fig. 3D) is higher in comparison to the serum condition. Overall, these analyses (Fig. 3C-D) reveal that under serum condition the promoter is maintained more often in the OFF state leading to higher level of fluctuations in the Nanog transcription than in the 2i condition.

### Model predicts the heterogeneous Nanog mRNA and protein expression patterns

Predicting the expression level of the Nanog mRNA and protein under different culture conditions is extremely important to reliably estimate the fluctuations present in the system. Keeping this in mind, we investigate the Nanog mRNA expression pattern under serum and 2i conditions, and found that the total Nanog mRNA expression shows a long tail distribution with a reasonably high variance under serum condition (Fig. 4A, upper panel) (Singer et al., 2014). This agrees with the experimental findings, where a shorter promoter ON state or a longer OFF state subsequently lead to long tail mRNA distributions with significant variances (Munsky, 2012; Nicolas et al., 2017). Subsequently, our simulations foretell that the mRNA distribution under 2i condition (Fig. 4A, lower panel) will have much less variability. How this kind of fluctuation pattern at the Nanog mRNA level affects the Nanog protein expression level under different culture conditions? To address this question, we estimated the Nanog protein expression level under different 2i conditions by comparing them to the serum condition (Fig. 4B). Interestingly, our model simulations predict that protein expression will increase under different 2i conditions in comparison to serum condition, which reconciles the experimentally observed Nanog protein expression pattern (Abranches et al., 2013).

This suggests that there may exist a favorable correlation between total Nanog mRNA and total Nanog protein in both serum and 2i culture conditions. Our model simulations confirm the existence of such correlated expression pattern of Nanog mRNA and protein under serum (Fig. 4C) and 2i condition (Fig. 4D), which had been observed experimentally by Ochiai et al. in ESC population. Collectively, this reflects that the expression variability in Nanog protein level mainly originates from the Nanog mRNA heterogeneity.

**Fig. 4.**
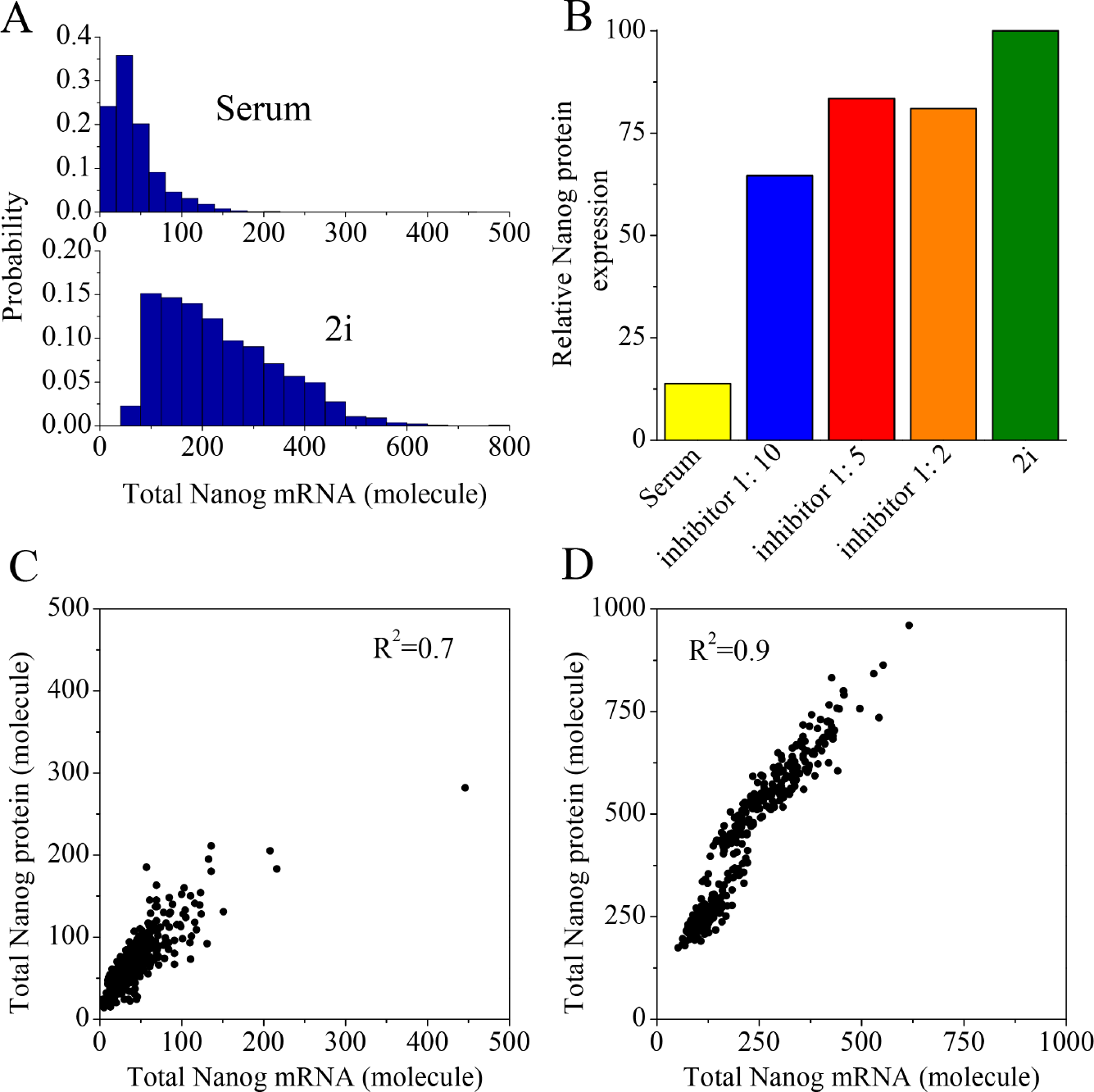
Simulated expression patterns of Nanog mRNA and protein under different culture conditions. **(A)** Distribution of mean Nanog mRNA transcripts per cells in serum (CV=77.5%) and 2i (CV=49%) culture conditions, respectively. **(B)** Nanog protein expression pattern under serum and different 2i inhibitor conditions (relative to the highest protein expression under normal (PD0325901: CHIR99021 = 1: 3) 2i condition). Correlation between Nanog mRNA and protein expressions under **(C)** serum and **(D)** 2i condition (considering 400 ESC’s).

### Model reconciles the robust noise partitioning in Nanog transcription

Being convinced that our model simulations were producing right kind of expression patterns of Nanog mRNA and protein, we employed stochastic simulations to verify whether our model can mimic the experimentally observed strict noise partitioning in Nanog transcriptional process or not. Our model simulations (Fig. 5A) indeed faithfully reproduced (Table 1) the ~ 45:55 ratio of intrinsic and extrinsic fluctuations in Nanog transcription under serum as well as in different inhibitor conditions (i.e., only MEK inhibitor (PD325901) and different 2i conditions), which is in accordance with the experimental observation. Not only that, we numerically simulated several other combinations of 2i conditions (Fig. 5B, Table 1), and found that the intrinsic and extrinsic fluctuations fascinatingly maintain the ~ 45:55 ratio.

**Fig. 5.**
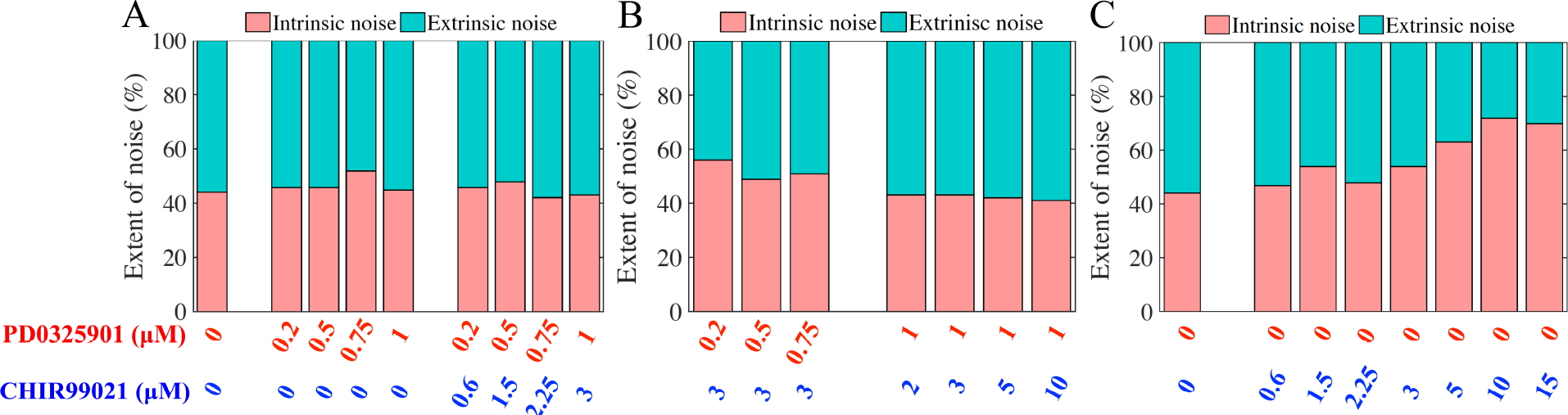
Model predicted % contribution of intrinsic and extrinsic noises under serum and different inhibitor conditions. **(A)** Stochastic simulations reconcile the experimentally observed (Ochiai et al., 2014) intrinsic and extrinsic noise ratio under serum, only PD0325901 inhibitor and specific 2i inhibitor doses. **(B)** Model simulations predict a sustain maintenance of ~ 45:55 ratio of intrinsic and extrinsic noise under other 2i conditions. **(C)** Model simulations predicted ratio of intrinsic and extrinsic noises under only CHIR99021 mediated inhibition, where higher doses of CHIR99021 will produce a higher level of intrinsic fluctuations.

Next, we computationally explored how to bypass this kind of strict intrinsic and extrinsic noise ratio maintenance? Importantly, our stochastic simulations (Fig. 5C, Table 1) unveil that by maintaining a relatively high level (10-15 *μ*M) of only CHIR99021 inhibitor (GSK3 inhibitor) condition will produce a much higher level of intrinsic (~ 70%) fluctuation in the system compared to extrinsic (~ 30%) noise. This prediction can be easily validated by making a simple smFISH kind of experiment performed by Ochiai et al. in this context with only CHIR99021 inhibitor.

### Dynamical reorganization of transcriptional events leads to precise apportioning of Nanog transcriptional fluctuations

At this juncture, we investigated the reasons behind such strict apportioning of the transcriptional fluctuations under such diverse inhibitory conditions, and why only employing higher level of CHIR99021 inhibitor circumvents such ratio maintenance? We used our model in a systematic manner to address the abovementioned question. It is well-known in literature that increase in either promoter activation rate or decrease in promoter deactivation rate lead to reduced intrinsic fluctuations, whereas increase in transcription rate elevates the intrinsic noise level (Fig. 1) (Ozbudak et al., 2002; Swain et al., 2002; Thattai and van Oudenaarden, 2001). Thus, we focused on how different inhibitory conditions are affecting rates (forward and backward) that control the promoter ON state maintenance, and the subsequent transcription rate to produce the Nanog transcripts (Supplementary file 4). Figure supplement 4 reveals that the application of only PD325901 inhibitor systematically increases both the promoter activation rate (without affecting deactivation rate), and the transcription rate (Fig. 1B), which ultimately increases the number of Nanog transcripts in the system (Table 1). This causes a systematic drop in the intrinsic as well as in the overall noise in a proportionate manner to maintain the ~ 45:55 noise ratio maintenance as a function of increasing doses of PD325901 (Fig. 6A).

Next, we investigated the effect of administrating the CHIR99021 inhibitor alone to the ESC’s culture medium. Figure supplement 4 demonstrates that till 3*μ*M dose of CHIR99021, the promoter activation and deactivation rates are marginally affected, while the transcription rate decreased gradually. All these lead to decrease in the intrinsic as well as the overall noise (Table 1) till 3*μ*M dose of CHIR99021, and a ~ 45:55 ratio is still maintained (Fig. 6B). With increasing doses of CHIR99021, the deactivation rate of the promoter gets highly affected (Figure supplement 4), which keeps the promoter mostly in the OFF state leading to a sudden increase in the intrinsic fluctuations (Fig. 6B). However, the overall absolute noise level remains relatively unaffected, which gets reflected in the decreasing level of extrinsic noise. This is how higher doses of CHIR99021 avoids the ~ 45:55 noise ratio maintenance.

We further analyzed the noise statistics under 2i conditions. Figure supplement 4 displays that under different combinatorial doses of PD325901 and CHIR99021 inhibitors, the increase in promoter activation rate and the subsequent decrease in both promoter deactivation and transcription rates reduce the intrinsic as well as the overall noise in Nanog transcription by increasing the Nanog transcripts level under these conditions (Table 1). To unveil which inhibitor plays a bigger role in Nanog transcriptional fluctuation, we analyzed and plotted our simulation results (Table 1) in different ways (Fig. 6C-E). Fig. 6C reflects the fact that even in the presence of high dose of CHIR99021 inhibitor (3 *μ*M), presence of PD325901 inhibitor takes control over the Nanog transcriptional noise regulation. This is true for any other 2i combination (Fig. 6D) doses, and importantly, even relatively higher doses of CHIR99021 inhibitor can not take control over noise regulation (Fig. 6E) in presence of moderately high dose of PD325901 inhibitor (1 *μ*M). All these results imply that PD325901 inhibitor plays a more influential role in governing Nanog transcriptional fluctuations, while CHIR99021 acts as a moderator for the same in presence of PD325901 inhibitor. However, our model predicts a greater role of CHIR99021 in fine tuning Nanog transcriptional noise in absence of PD325901 inhibition.

**Fig. 6.**
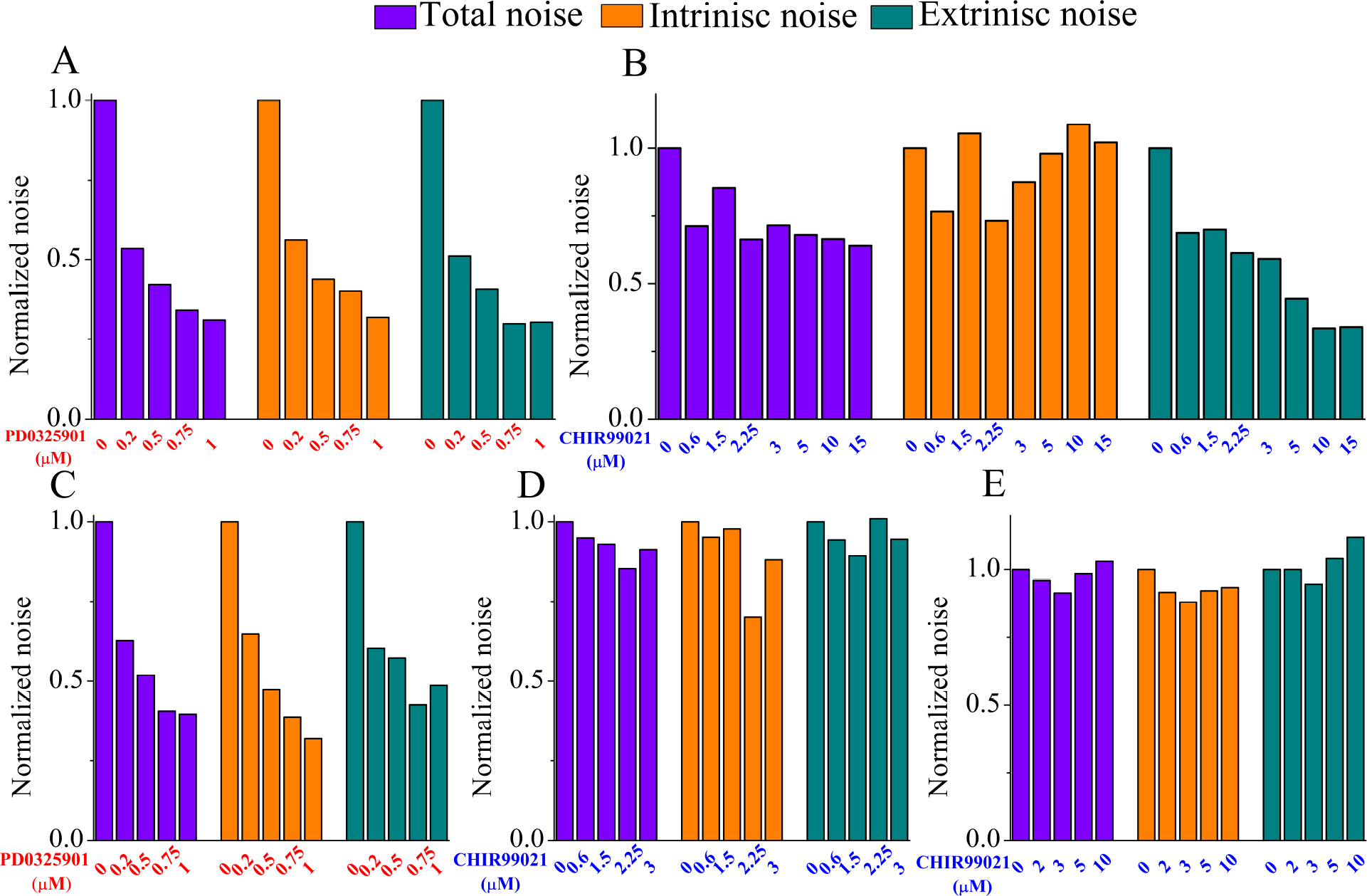
Analysis of the relative contributions of PD0325901 and CHIR99021 inhibitors on Nanog transcriptional regulation. Effect of increasing doses of only PD0325901 inhibitor **(A)**, and only CHIR99021 inhibitor **(B)** on total, intrinsic and extrinsic noises (in both the cases, fluctuations are relative to serum condition). (**C**) Effect of increasing doses of PD0325901 inhibitor in presence of constant 3 μM CHIR99021 inhibitor dose on total, intrinsic and extrinsic noises (relative to fluctuations for only 3 μM CHIR99021 inhibitor condition). (**D**) The variation in noises under different 2i inhibitor concentrations, where PD0325901 and CHIR99021 inhibitor concentrations maintain a ratio of 1:3 each time. For each column, the relative change in any noise under a respective 2i condition is calculated with respect to the same noise corresponding to only PD0325901 inhibitor condition. (**E**) Effect of increasing doses of CHIR99021 inhibitor in presence of constant 1 μM PD325901 inhibitor dose on total, intrinsic and extrinsic noises (relative to fluctuations for only 1 μM PD325901 inhibitor condition).

## Conclusion

Despite manifesting a highly heterogeneous expression pattern in ESC’s under different culture conditions, Nanog is known to be a key regulator of the cell fate decision making events in ESC’s. Thus, it is imperative to understand what causes this inherent heterogeneity in Nanog expression to direct the developmental transitions in ESC’s in an informed manner. It was believed that transcriptional fluctuations might play a crucial role in maintaining such heterogeneity. However, recent experiments demonstrated that the overall fluctuations at the Nanog transcriptional level gets uniquely partitioned with a ~ 45:55 ratio of intrinsic and extrinsic noises, under various culture conditions. By employing a new quantitative stochastic simulation (supplementary file 5) study of a simple Nanog regulatory network (Fig. 1), we explored what really controls this bizarre balance of Nanog transcriptional noise partitioning under various culture condition. Our proposed numerical framework (Fig. 2) allowed us to incorporate the effects of various extrinsic variability’s such as cell division, transcriptional rate variation dependent on the cell cycle phases, and many other such external factors that are often ignored in a conventional SSA simulation. Importantly, this method can be implemented in general for any regulatory network functional within a cell to qualitatively capture the sources of external variability’s without considering an explicit cell cycle network model.

Employing our numerical simulation method, we reproduced the expression patterns of Nanog mRNA and protein (Fig. 4), and qualitatively reconciled the ~ 45:55 ratio of intrinsic and extrinsic noises (Fig. 3) observed experimentally under different culture conditions (Table 1, Fig. 5). We must emphasize the fact that we have strictly followed experimental literature to phenomenologically introduce the effect of inhibitors in Nanog transcriptional regulation. Not only that, we have performed a systematic sensitivity analysis (Figure supplement 5), which exhibits that our simulation results are not much sensitive on the parameter values used. By analyzing our stochastic simulation results (Fig. 6), we found that this unique partitioning of noises under different culture condition is organized by balancing the rates of promoter activation/deactivation and the rate of transcription from the promoter ON state (Figure supplement 4). Our analyses uncover that PD0325901 inhibitor controls the overall fluctuations at a much greater extent (Fig. 6A), and even under 2i conditions (i.e., in presence of CHIR99021), the effect of it dominates over CHIR99021 activity (Fig. 6C-E). Only in absence of PD0325901 inhibitor, one can realize the true effect of CHIR99021 (Fig. 6B) on transcriptional fluctuations of Nanog. Importantly, our model predicts that at higher doses of CHIR99021 the ratio (~45:55) of intrinsic to extrinsic noises can be avoided due to relatively higher contributions of intrinsic noise.

In summary, our stochastic modeling study provides an improved way to understand the Nanog transcriptional heterogeneity maintenance under different culture conditions. We have shown how and why different inhibitors maintain a strict partitioning of transcriptional fluctuations in Nanog, and predicted avenues to experimentally evade such noisy regulation. Thus, our modeling study not only predicts new experiments, but put forwards new possibilities to control ESC dynamics in an improved manner. This was made possible by our newly proposed numerical set up (Fig. 2, Figure supplement 3), which we believe will find a wide applicability not only in the context of Nanog regulation, but in understanding many other relevant biological problems.

## Acknowledgements

Thanks are due to IIT Bombay for providing fellowship to TS. This work is supported by the funding agency DBT, India grant (BT/PR11932/BRB/10/1315/2014).

---

**Figure supplement 1.**
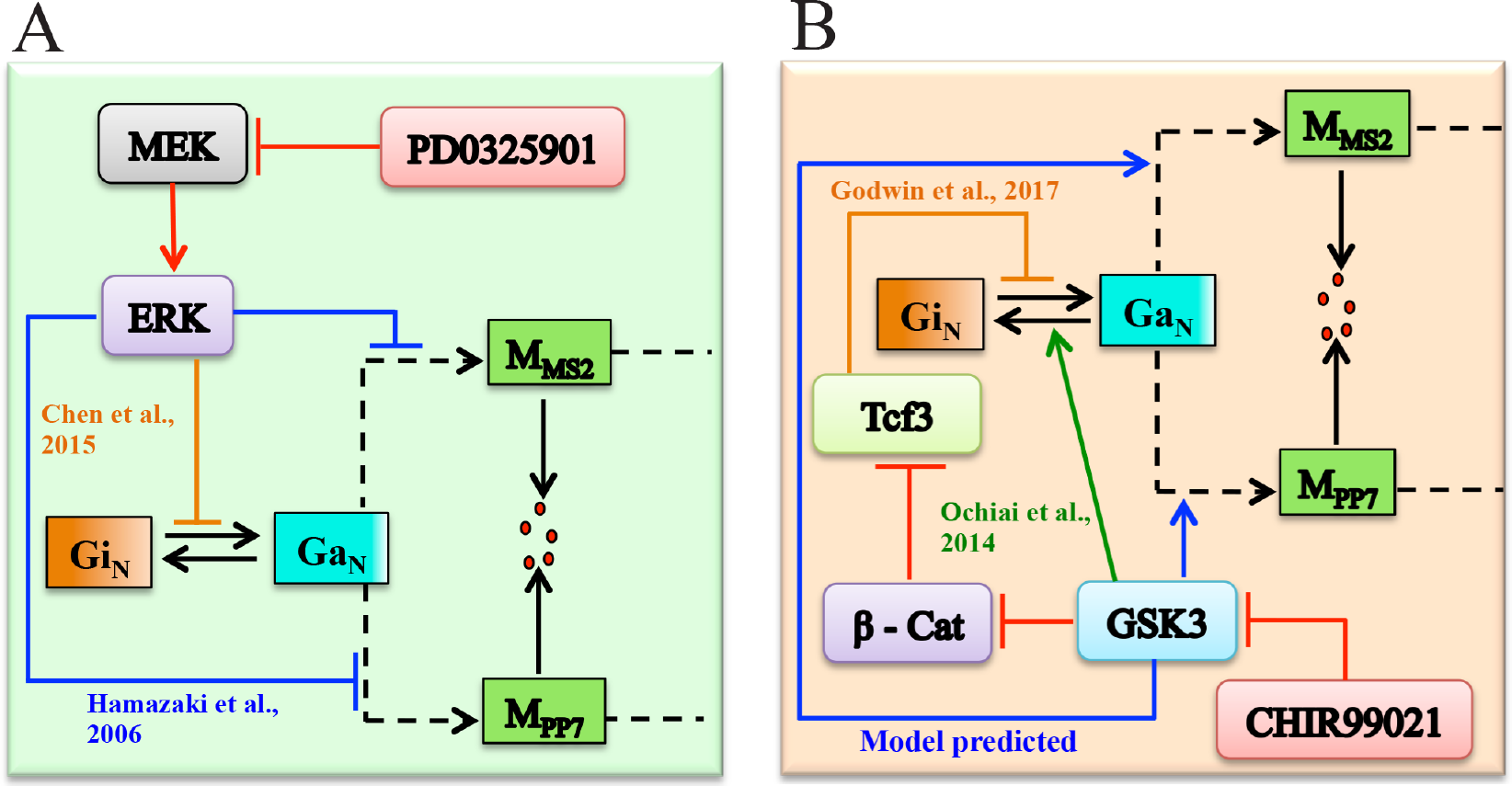
The detailed regulatory interaction of Inhibitors. **A**. The effect of PD0325901 inhibitor on Nanog gene activation and transcription rates through ERK signaling via MEK inhibition. **B**. Influence of CHIR99021 inhibitor on Nanog gene-activation and transcription rates through some intermediate steps.

**Figure supplement 2.**
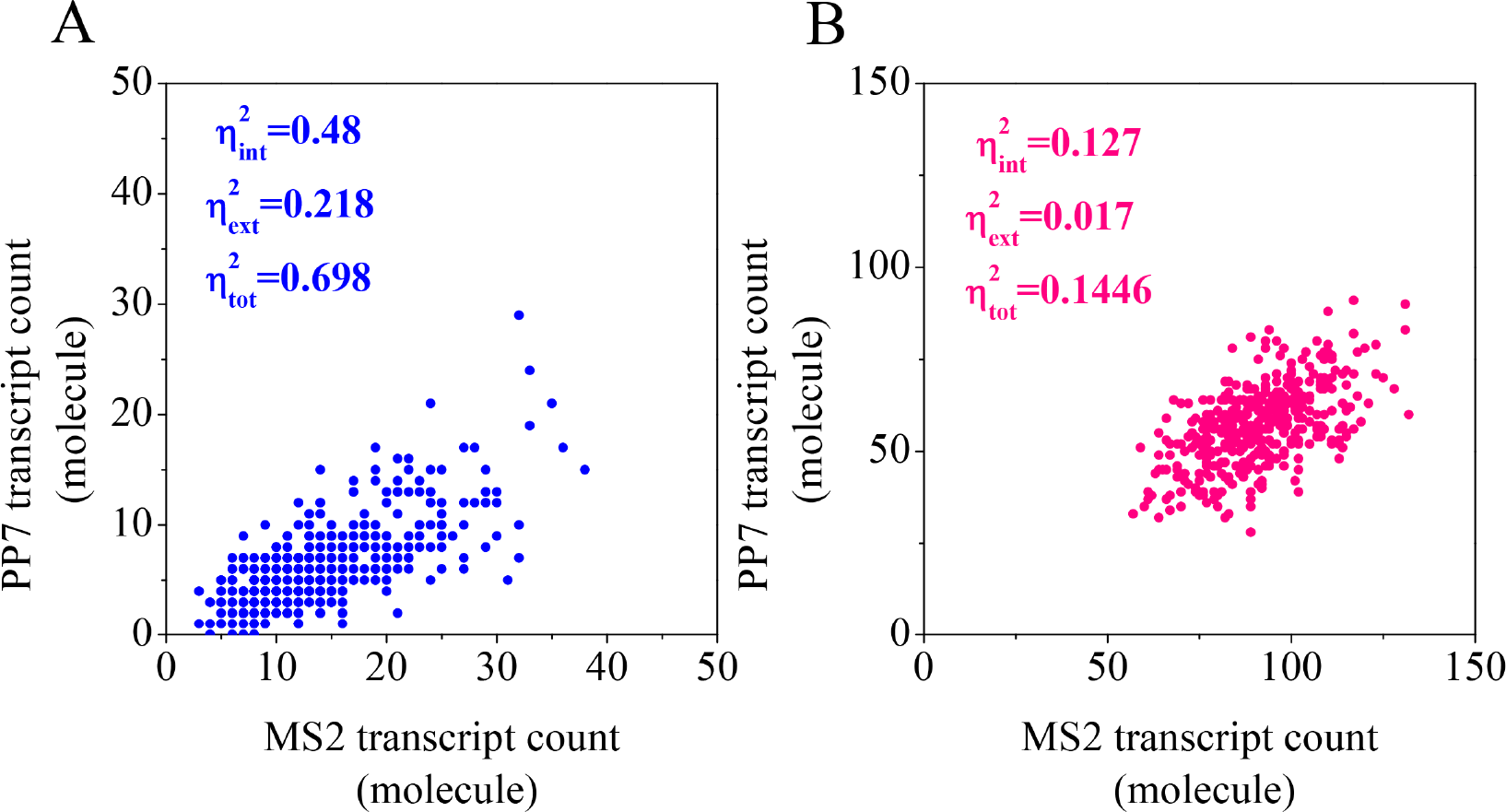
Intrinsic noise mediated Nanog expression variability without considering cell division as an extrinsic noise source. Scatter plot of Nanog mRNA transcripts in **A**. serum and **B**. 2i culture conditions. Each dot represents mean number of Nanog-MS2 and Nanog-PP7 transcript in a single cell. 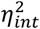, 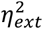 and 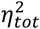 represent intrinsic, extrinsic and total noise respectively. The corresponding noise values are given into the figures respectively.

**Figure supplement 3.**
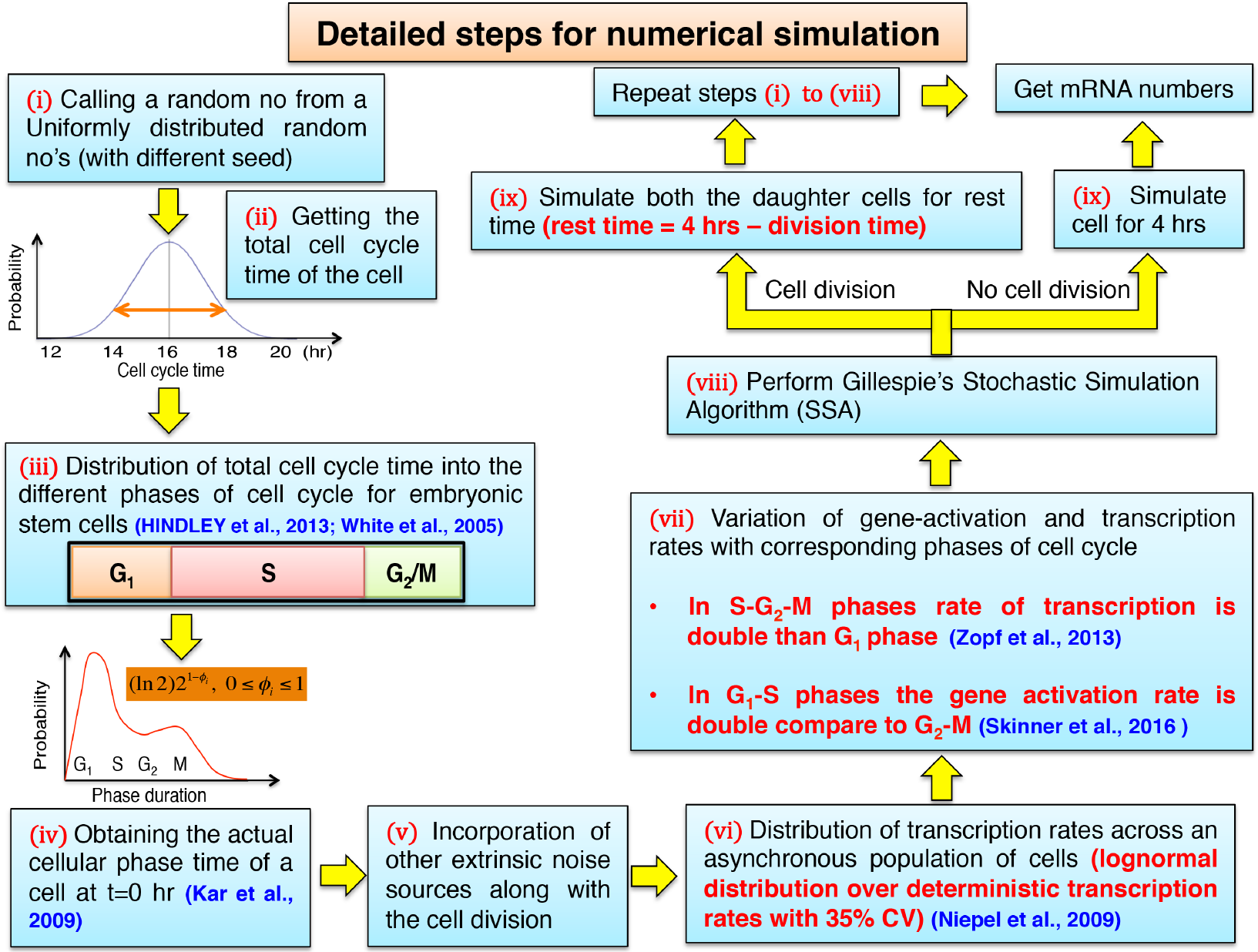
The detailed steps involved in the numerical simulation by employing SSA. Here we have incorporated other extrinsic noise sources along with the cell division.

**Figure supplement 4.**
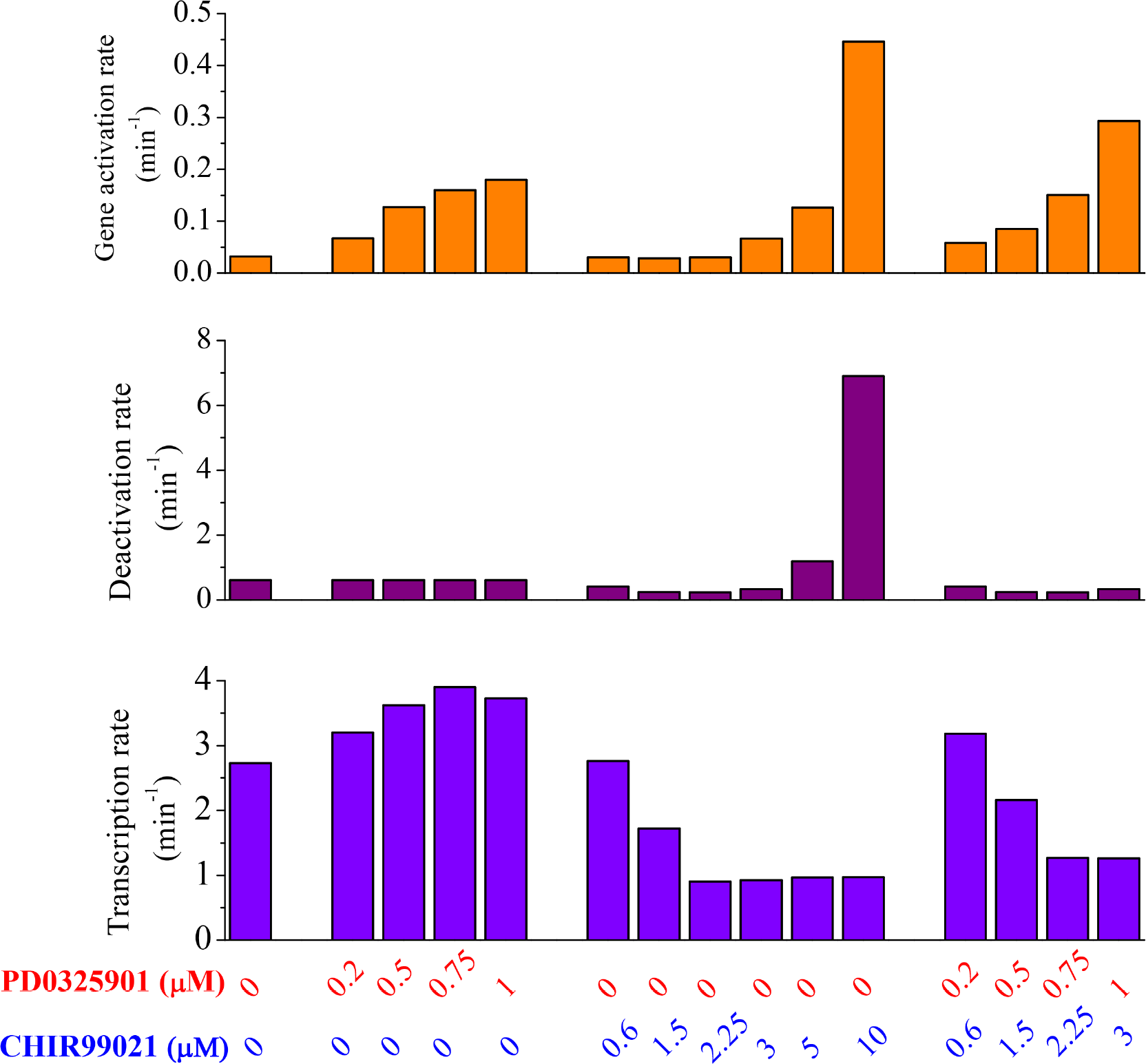
Distribution of rates under different culture conditions. (i) In presence of Serum (ii) In presence of 0.2 μM, 0.5 μM, 0.75 μM and 1 μM PD0325901 inhibitor concentration respectively (iii) In presence of 0.6 μM, 1.5 μM, 2.25 μM, 3 μM, 5 μM and 10 μM of CHIR99021 inhibitor concentration respectively. (iv) Under different 2i culture conditions maintaining the concentration ratio of PD0325901 and CHIR99021 inhibitors as 0.2:0.6, 0.5:1.5, 0.75:2.25 and 1:3 respectively. **A.** Distribution of gene activation rate under different culture conditions. We have calculated the rates by considering the positive feedback coming from oct4-sox2 heterodimer complex. **B.** Distribution of deactivation rate under the above mentioned culture conditions. **C.** Distribution of transcription rate for Nanog-MS2 transcript under different culture conditions.

**Figure supplement 5.**
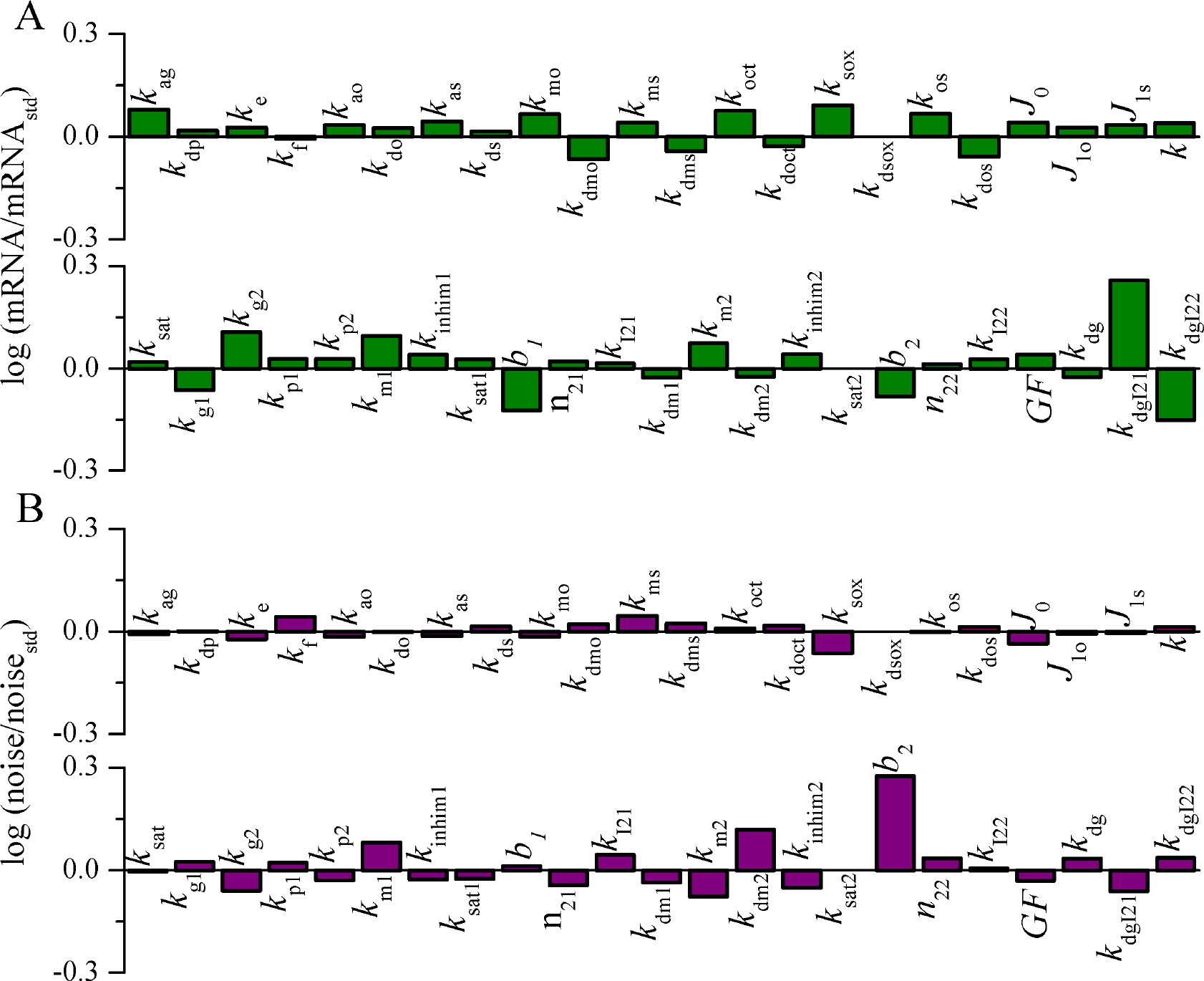
Sensitivity analysis of the parameters involved in the model. In all the cases, parameters are increased individually (about 20% of the values provided in the Table S2) keeping all other parameters constant. **A.** The change in mRNA number is calculated on the basis of change in the number of mRNA observed with respect to standard amount of mRNA under 2i condition (PD0325901and CHIR99021 inhibitors concentration are 1 and 3 μM respectively). We have stochastically calculated the number of mRNA for 400 cells over 4 hours in presence of all noise sources. **B**. The sensitivity is measured by measuring the changes in total noise as one of the parameter is increased individually about 20% of its original value (provided in Table S2), and the total noise is compared with respect to the total noise under 2i condition (PD0325901and CHIR99021 inhibitors concentration are 1 and 3 μM respectively).

